## Supplemental Information for "The flexible stalk domain of sTREM2 modulates its interactions with brain-based phospholipids"

Boulder, CO, USA

^2^Department of Endocrinology, Metabolism, and Diabetes, University of Colorado Anschutz Medical Campus, Aurora, CO, USA

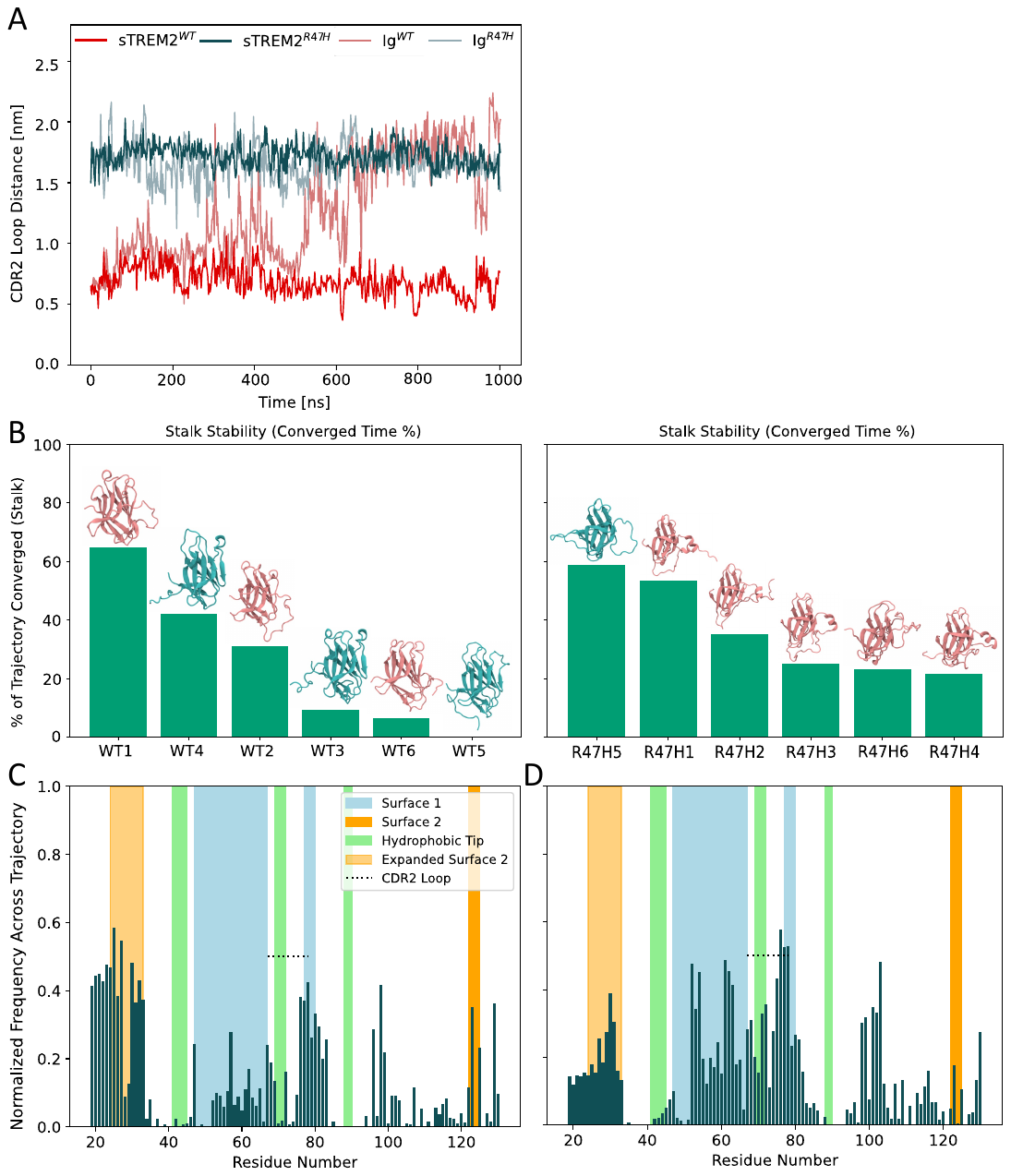

**Figure S1.** (A) Distance between CDR loop residues 45 and 70 over time for WT and R47H models of sTREM2 and TREM2 (Ig). (B) Percentage of simulation time that the WT (left) and R47H (right) sTREM2 stalk region remained within a converged RMSD window (±2 Å over 50 ns) for each replicate, calculated using MDAnalysis, shown alongside a visualization of a representative structure from each simulation, color-coded by stalk position (pink for ‘Surface 1’ and blue for ‘Surface 2’). Normalized fractional occupancy of residues in the Ig-like domain of (C) sTREM2^WT^ and (D) sTREM2^R47H^ by the stalk, averaged across six replicates each.

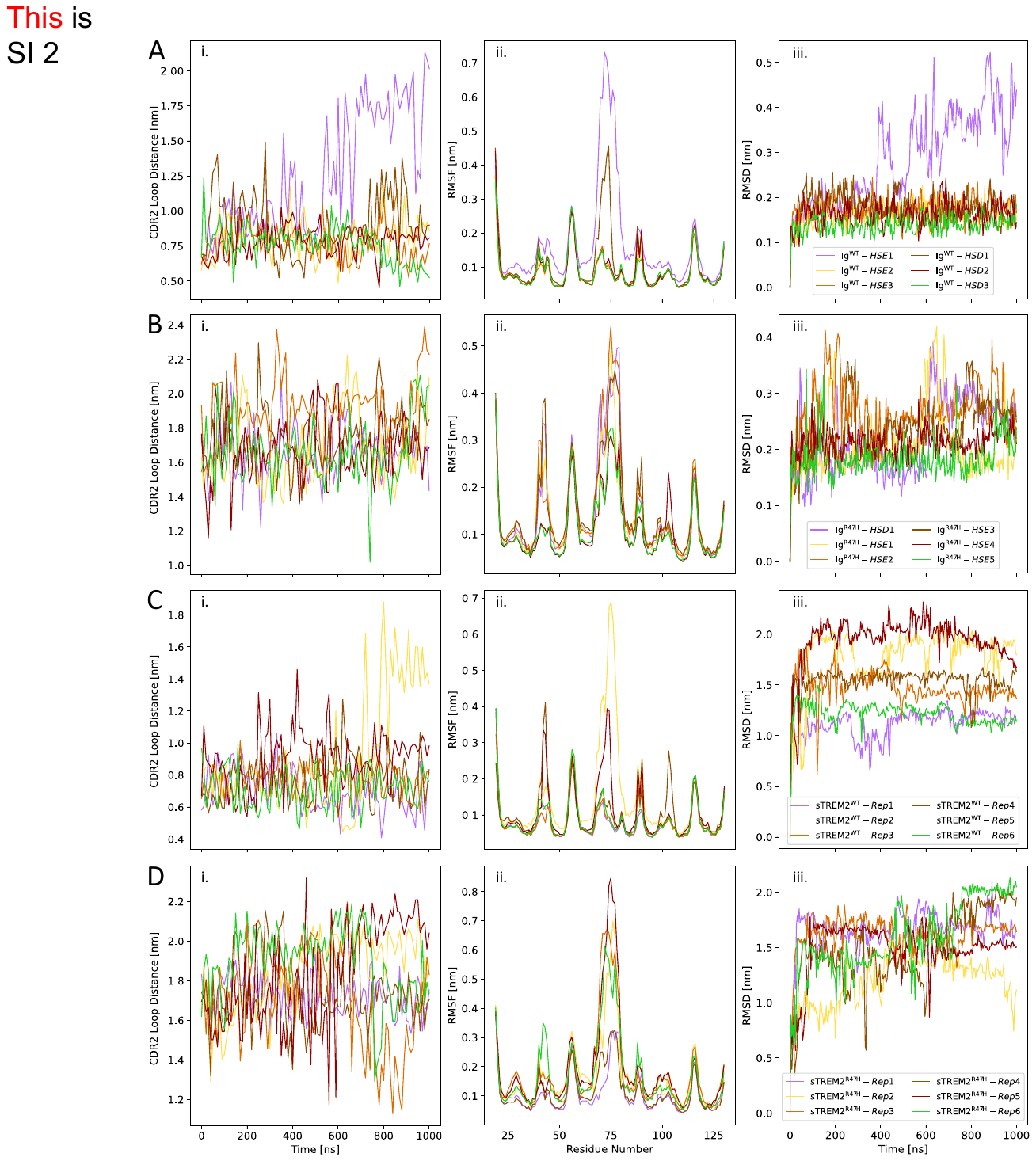

**Figure S2.** Analyses across MD replicates for (A) Ig^WT^, (B) Ig^R47H^, (C) sTREM2^WT^, and (D) sTREM2^R47H^. Each row shows: (i) distance between CDR loop residues 45 and 70 versus simulation time, (ii) time-averaged Cα RMSF per residue, and (iii) Cα RMSD over time.

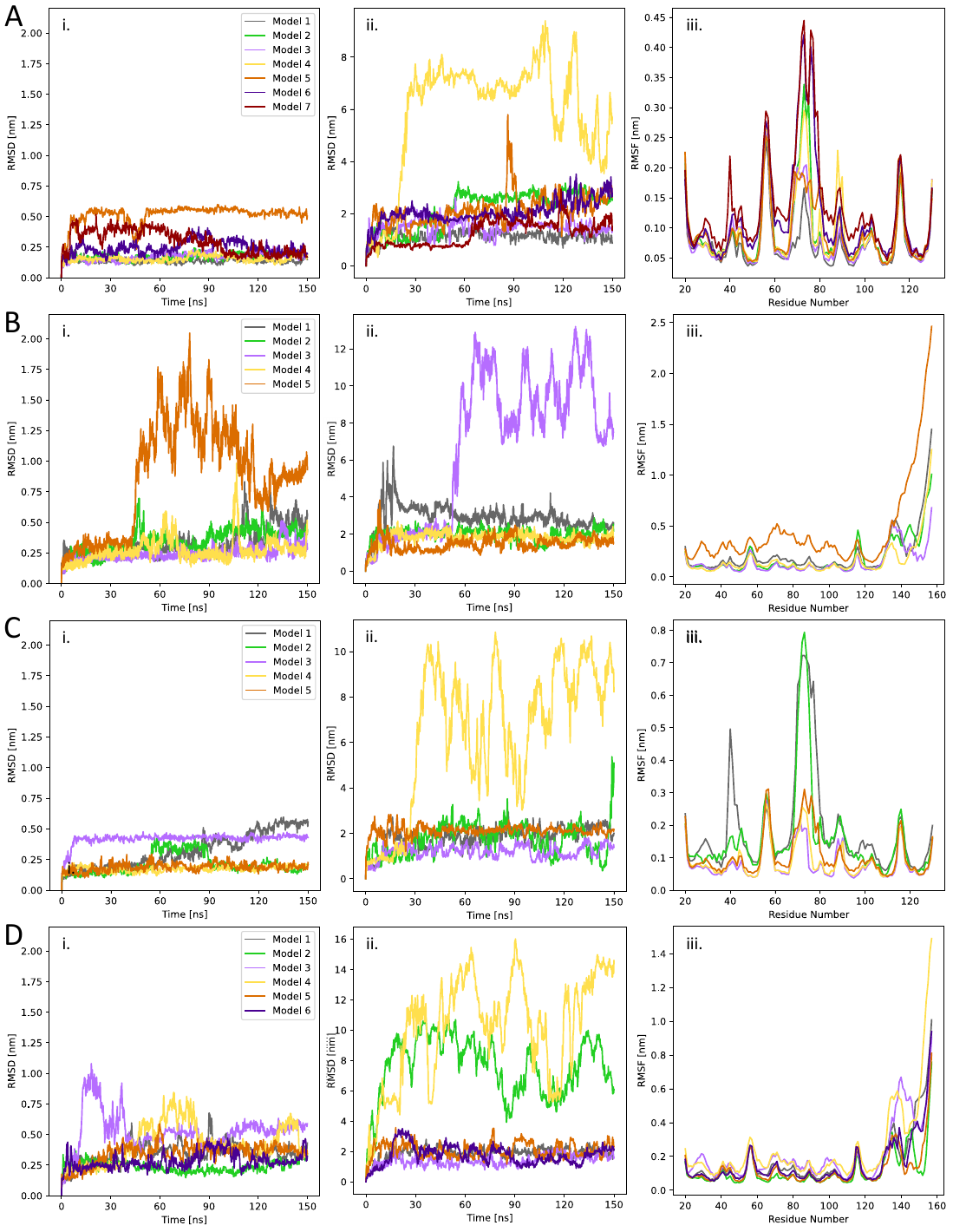

**Figure S3.** Comparative RMSD and RMSF analyses across individual MD trials for each WT (s)TREM2/PL system: (row A) SOPS/Ig^WT^, (row B) SOPS/sTREM2^WT^, (row C) SOPC/Ig^WT^, and (row D) SOPC/sTREM2^WT^. The subfigures in each row show: (i) Cα RMSD of the protein structure vs. simulation time, (ii) RMSD of the PL calculated relative to the protein vs. simulation time to capture broad PL movements, and (iii) temporally averaged Cα RMSF of protein residues.

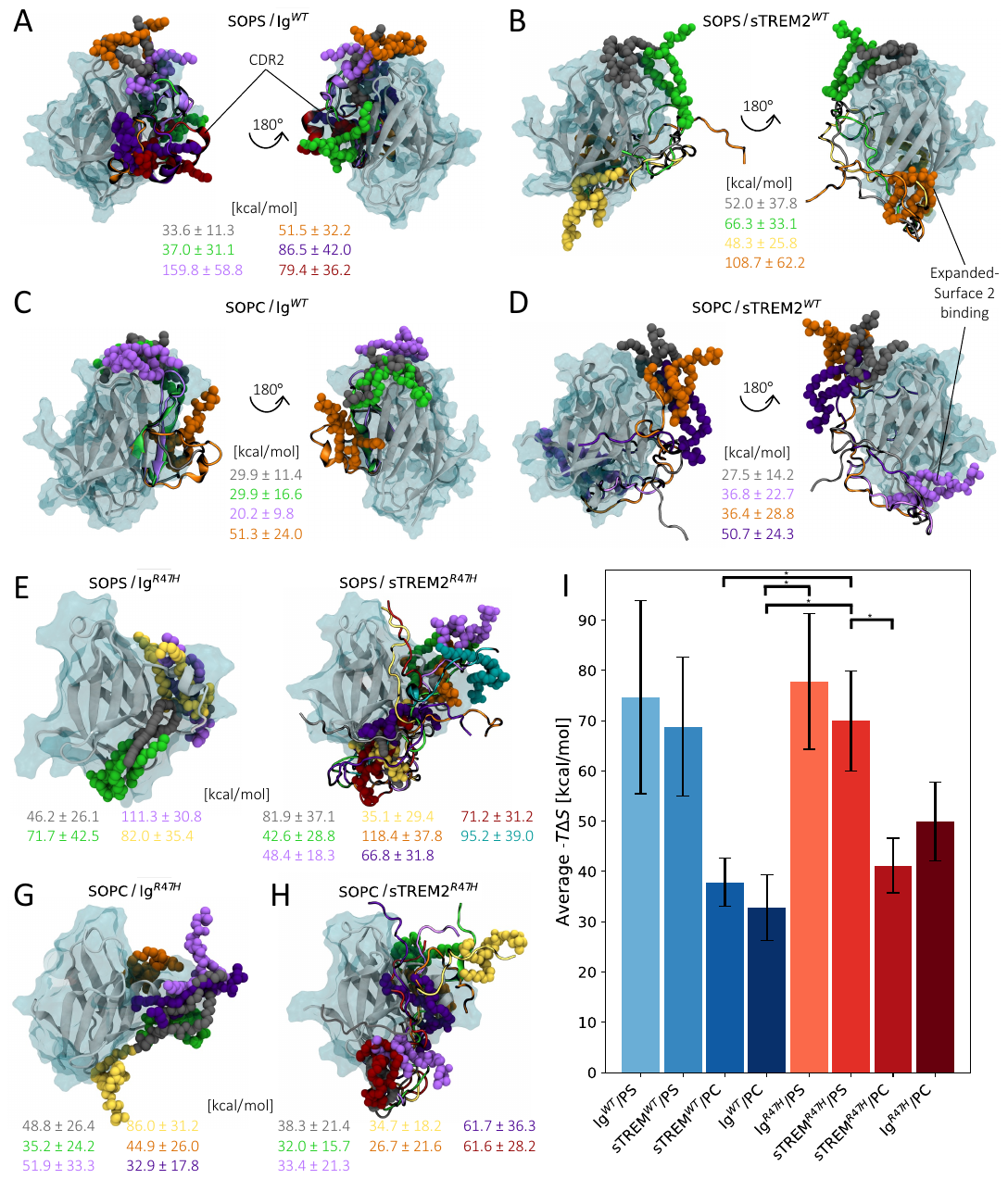

**Figure S4.** VMD-rendered images from PL/(s)TREM2 simulations (A-H), with the PLs colored to match their binding interaction entropy contributions (-TΔS) (kJ/mol) in panel I. (I) Comparison of the interaction entropy contributions averaged across all WT and R47H models for each PL/protein system. Errors bars represent the standard error of the mean. Black brackets with a single asterisk represent statistical significance with 0.01<p<0.05.

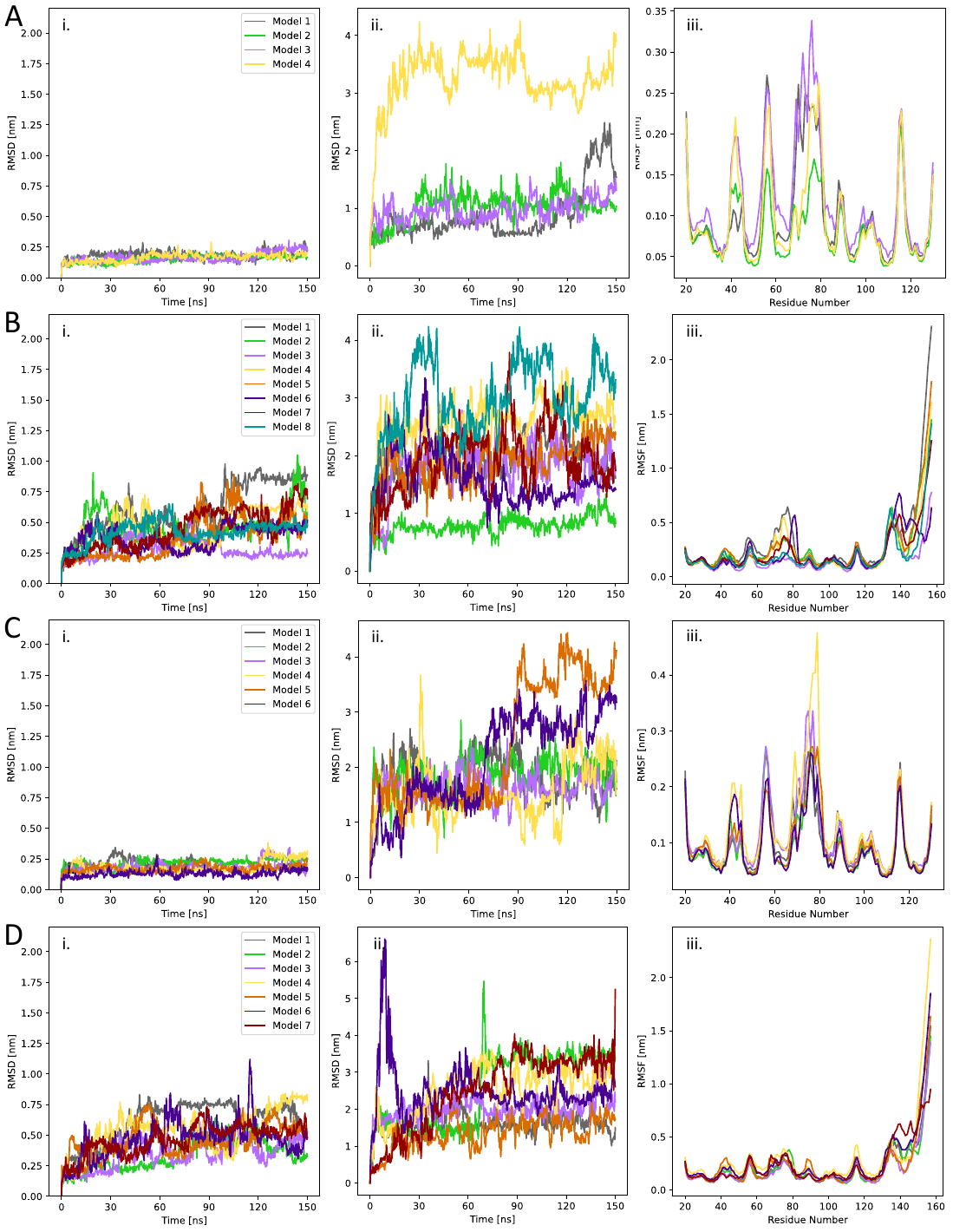

**Figure S5.** Comparative RMSD and RMSF analyses across individual MD simulation trials for each R47H (s)TREM2/PL system: (row A) SOPS/sTREM2^R47H^, (row B) SOPS/Ig^R47H^, (row C) SOPC/sTREM2^R47H^, and (row D) SOPC/Ig^R47H^. The subfigures in each row show: (i) Cα RMSD of the protein structure vs. simulation time, (ii) RMSD of the PL calculated relative to the protein vs. simulation time, and (iii) temporally averaged Cα RMSF of protein residues.
